# Can lateral tenodesis improve the rotational stability of the ACL reconstruction? A finite element analysis

**DOI:** 10.1101/2023.10.08.561440

**Authors:** Konstantinos Risvas, Dimitar Stanev, Konstantinos Moustakas

## Abstract

One of the most common knee injuries is the Anterior Cruciate Ligament (ACL) rupture with severe implications on knee stability. The usual treatment is the ACL Reconstruction (ACLR) surgery where the surgeon replaces the torn ligament with a graft in an effort to restore knee kinematics. In case of excessive rotatory instability, Lateral Extra - Articular Tenodesis (LET) can be performed in combination with ACLR. Additionally, LET appears to reduce ACLR graft forces reducing graft failure chances. However, there are concerns about overconstraining physiological rotation. To gain insight in this controversial topic, we developed an automatic, open-source tool to create a series of Finite Element (FE) models attempting to investigate the interactions of ACLR and LET through simulation. We started by creating a validated model of the healthy knee joint that served as reference for subsequent FE simulations. Then, we created FE models of standalone ACLR and combined ACLR - LET. Each model was assessed by applying a loading profile that resembles the Pivot - Shift clinical exam. We measured the External Tibia Rotation (ETR), the Anterior Tibia Translation (ATT) of the lateral tibial compartment, and the ACLR graft stress developed around the femoral tunnel insertion site. We observed the following: a) LET reduces ETR and ATT compared to isolated ACLR, b) combined ACLR - LET is more sensitive to LET graft pretension with lower values showcasing performance closer to the healthy joint, c) LET reduces ACLR graft forces for the same pretension values, d) LET exhibits significant overconstraint for higher pretension values. In general, these findings are in agreement with relevant clinical studies and accentuate the potential of the developed framework as a tool that can assist orthopaedists during surgery planning. We provide open access for the FE models of this study to enhance research transparency, reproducibility and extensibility.

## Introduction

Knee joint injuries have become a commonplace in demanding physical activities and competitive sports, such as soccer, basketball and skiing [1–3]. One of them is the typical Anterior Cruciate Ligament (ACL) injury where the Native - ACL ligament is partially or totally ruptured. The ACL’s primary biomechanical role is to restrain Anterior Tibial Translation (ATT). It also acts as a secondary rotational stabilizer, reducing excessive External Tibial Rotation (ETR). Consequently, such an injury reduces the physical capability and deteriorates the quality of life of the injured population with further implications on their psychological condition. Moreover, ACL demonstrates poor healing capability which is drastically eliminated in total rupture. Therefore, a treatment to restore knee kinematics is of paramount importance, especially in the case of professional athletes.

The type of treatment is usually selected based on the long term objectives and whether the injured person wishes to resume demanding physical activities in the near future. Assuming this requirement, a surgical treatment to the injury is the standard approach, and the Anterior Cruciate Ligament Reconstruction (ACLR) has emerged as the prevalent surgical method. Describing ACLR in a nutshell, the surgeon clears the remnants of the torn native tissue, drills two tunnels in the proximal tibia and distal femur based on the anatomical ACL footprints, and places through the tunnels a replacement graft [4, 5]. Usually, this graft consists of tissue extracted from a harvesting site of the same patient (autograft) or another donor (allograft). Apart from graft tissue, ACLR is a demanding surgical procedure where a plethora of additional parameters needs to be determined. The adopted number of osseous tunnels and their orientation through each bone leads to different techniques, such as Single Bundle (SB) and Double Bundle (DB) [6–11]. Moreover, the graft tension and knee angle prior and during graft fixation are important ACLR features [9, 12].

Although ACLR exhibits excellent functional restoration of ATT, the reestablishment of rotational knee stability is still questionable. This type of instability is also related to ACLR graft rupture and early progression of Osteoarthritis (OA) [13, 14]. Towards this direction, additional surgery techniques that are often performed along ACLR have been developed. Lateral Extra-Articular Tenodesis (LET) is a renascent surgical approach that aims to enhance knee rotational stability and is usually performed along ACLR especially on athletes that perform activities where cutting and pivoting are dominant [13–16]. From a clinical perspective, LET is performed by drilling a tunnel posterior and superior to the Lateral Collateral Ligament (LCL) femoral attachment. The graft is usually a 10mm wide strand stripped of the Iliotibial Band (ITB) [14, 17, 18]. The surgeon passes the graft beneath the LCL, applies a pretension load and fixes it at a given knee flexion angle. Apart from the specific surgical parameters and techniques, the surgeon’s ability and experience are critical factors that affect the outcomes of ACLR. Also, the optimal combination of the surgery-related criteria for each individual patient is not known in a pre-surgical setting. Rather, the ACLR surgical results are evaluated when each subject resumes physical activities which impose high loads to the reconstructed knee joint. An *in vivo* evaluation of knee kinematics and kinetics after ACLR is a very demanding task that requires the conduction of expensive and complex experiments [19]. Clinicians usually resort to clinical exams such as the Lachman and Pivot - Shift (PS) tests for surgery assessment. The former performs outstandingly well when assessing ATT [20]. On the other hand, the latter is better suited for assessing anterolateral knee instability. However, these tests depend on the clinician’s experience and feature high sensitivity and specificity [19–22]. An alternative approach is to deploy robotic simulators that are programmed to apply suitable loading profiles in an effort to eliminate the subjective assessment of knee stability [23].

All these pitfalls have led to the emerge of biomechanics simulations as a valuable alternative tool, available to clinicians, engineers and researchers. These simulations are based on computational numerical techniques with the Finite Element (FE) method being one of the most prominent. In the case of ACLR, several FE studies can be found in literature. A common starting point is the creation of a subject-specific FE model [24] which is validated in terms of joint kinematics and material properties. Initially, Magnetic Resonance Imaging (MRI) and computational tomography (CT) data are segmented and reconstructed to acquire a boundary geometrical representation of the underlying subject-specific anatomical structures. Then, the obtained surface meshes are usually filled with quadrilateral (quad) or hexahedral (hex) elements and are assigned realistic mathematical material models. ACLR is represented by excluding the native tissue and creating graft models. Grafts and the other knee ligaments can be modeled as 1D elements, such as springs that feature tension-only force - displacement behavior [25–30]. Alternatively, they have been also modeled as 3D volumetric meshes. The ligament meshes are usually reconstructed from MRI data, whereas the grafts are generated as cylindrical-shaped objects using dedicated software [30–41]. The material properties assigned to these geometries are usually adopted from cadaver studies. The graft models are adjusted to represent various harvesting areas [30–32, 36, 37, 40, 41]. The menisci and cartilages are usually modeled as 3D meshes with suitable material properties [24, 25, 42–44]. Furthermore, knee joint kinematics are modeled by imposing suitable rigid body constraints that are adjusted based on *in vivo* or *in vitro* experiments where the knee joint is mechanically tested [27–29, 45, 46]. These validated models also serve as reference points for subsequent researches [30, 31, 33, 34, 36].

Regarding boundary conditions, most of the FE ACLR studies simulated clinical exams, such as the Lachman Test (or Anterior Drawer test) [27, 28, 31–33, 37–40, 47] and the PS test [37, 40, 48–50]. Simulations of dynamic physical activities, such as gait can be also found [29, 47]. The results are usually knee joint kinematics or internal forces (stresses) subject to the adopted material model and compared to clinical data for model validation. Many studies evaluated knee kinematics restoration and graft loading based on anatomical tunnel placement [29, 34, 36, 39, 40, 47, 51–56]. Other ACLR studies focused on assessing the effect of other ACLR parameters, such as graft dimensions, graft pretension and graft fixation angle [30–32, 38]. On the other hand, to our knowledge the literature lacks systematic FE studies that attempt to model and assess the combined ACLR - LET surgery. As we mentioned, LET is performed when the ultimate objective is to restore knee rotational stability, albeit caution is required to avoid excessive rotational constraint, a situation that can lead to increased knee stiffness and loading to the lateral knee structures [57]. Additionally, weakening of the ITB due to graft harvest can further lead to knee loading implications due to movement alterations during dynamic activities [58]. These concerns served as motivation for us to develop a FE study to investigate the presumed advantages and drawbacks of the combined ACLR and LET surgery.

Towards this direction, we expand our previously published workflow [30] to encompass LET modeling. As before, the entire pipeline operates with open source software. The FE model assembly is performed through a custom Python module that is available in public. We have created a validated FE model that represents the healthy knee joint and serves as the baseline for ensuing ACLR - LET simulations. These include variations of the reconstructed knee in terms of adopted surgery (standalone ACLR or combined ACLR - LET) and graft pretension. We also have adopted a loading profile that is capable to produce the PS effects in a simulation setting. We use this profile to assess ETR and ATT restoration and graft stresses after ACLR reconstruction. Our findings suggest that the combined ACLR LET leads to reduced ETR compared to the standalone ACLR. It also causes a modest reduction of the ACLR graft stresses. An interesting finding is that the ETR is more sensitive to the LET graft pretension. On the other hand, the ATT depends on both the ACLR and LET graft pretension values. Moreover, larger LET pretension values may lead to excessive knee constraint on both ETR and ATT. To the best of our knowledge, this is the first FE simulation that attempts to investigate the effect of LET in restoring knee rotational stability. We have compared our results with clinically relevant studies. Finally, we share all created FE models with the research community aiming towards reproducibility, extensibility and enhanced research translucence.

## Materials and methods

In Fig 1, we present an overview of the adopted methodology and the distinct building blocks of the proposed workflow. In summary, we utilize MRI data acquired form the OpenKnee (s) project [59] and segment them to acquire surface representations of the anatomical structures. Subsequently, we employ open - source custom tools to create suitable volumetric meshes for the ligaments, cartilages, and menisci [60]. Additionally, we have created the “Surgery Modeling” tool to model the ACLR surgery procedure. This tool utilizes Blender scripting, to create the drilled bones and graft geometries. The cornerstone of this work is the “FEBio Exporter” tool, which we utilize as an interface to build all FE models, and execute them with the FEBio solver [61]. Moreover, we have developed scripts for experimental curve-fitting to estimate material parameters for the selected grafts. Also, We have experimented with boundary conditions representing the PS clinical exam and applied those to subsequent FE simulations to investigate the performance of the various ACLR choices.

**Fig 1.**
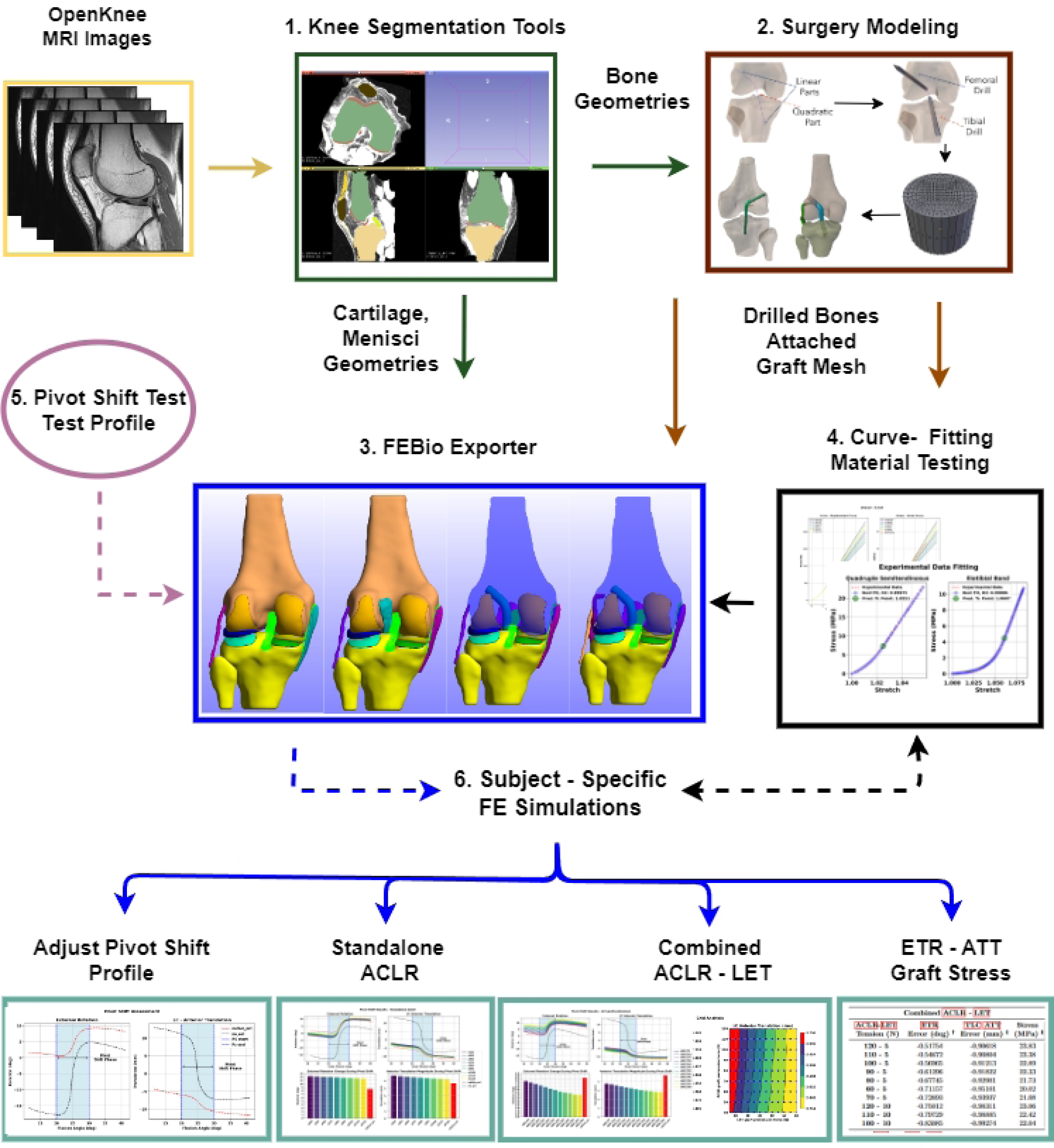
Overview of the proposed workflow. 1. The workflow starts with segmentation of MRI data to obtain subject specific anatomical geometries. 2. We developed the “Surgery Modeling” tool that utilizes Blender scripting to model the ACLR parts (drilled bones and grafts). 3. We created the “FEBio Exporter” tool to assembly subject - specific FE models of the healthy, injured and ACLR knee and to solve them using the FEBio software. 4. We adjusted graft material properties based on experimental data. 5. We developed a suitable loading profile in an effort to simulate the standard physical clinical exam of PS. 6. We performed a series of FE simulations to compare the standalone ACLR versus the combined ACLR - LET surgery techniques in terms of kinematics restoration and ACLR graft stress development.

### Surgery modeling

In this section, we will describe the developed “Surgery Modeling” tool that can be used to create ACLR knee models, suitable for FE analysis, similar to our previous work [30]. The tool accepts as inputs the segmented and reconstructed bone geometries and various user-defined parameters related to ACLR surgery. These include the adopted surgery approach, the corresponding bone surface landmarks that feature as entry and exit points of the bone tunnels, and the graft geometric characteristics, such as radius, mesh density and volumetric mesh type (hexahedral or tetrahedral). Using the desired landmarks as control points, we create a spline curve that acts as the surgery planning trajectory. Then, we create cylinder objects that act as drills, that are properly attached along the curve. By applying Boolean modifiers we create the tibial and femoral tunnels. We also utilize the curve to attach the created graft geometries between the tunnels. Finally, different face and element sets of the bone and graft meshes are stored for automatically applying proper boundary conditions during subsequent FE simulations.

In this work, we utilized this tool to model different versions of ACLR procedures, namely the standalone SB ACLR and the combined ACLR - LET surgeries. The bones were acquired by the OpenKnee (s) database and correspond to the subject “oks003”. The SB ACLR graft features a radius of 4mm [9]. The ITB graft is 10 mm wide close to its attachment to the tibia bone to resemble the ITB strip in relevant clinical surgeries [14, 17, 18]. A femoral tunnel with diameter of about 5 mm was created for graft fixation. Using the trajectory planning curve the graft was passed superior to the proximal LCL and through the femoral tunnel. The graft distal part was considered fixed with the tibia. We should emphasize that a segmented geometry of the ITB was not available, thus, we decided to place the proximal ITB end close to the Gerdy’s tubercle area of the tibia bone [14, 62].

### FE model assembly

In this section, we describe the workflow for creating the FE models. For each simulation scenario we describe the used geometries, adopted materials, enforced contact models and applied boundary conditions. We have developed the “FEBio Exporter” tool (Fig 1) for automated creation of FE models suitable for the FEbio solver. This provides us the ability to create multiple instances of models, each featuring different properties. Therefore, these models can be efficiently deployed in sensitivity analyses to investigate what-if scenarios, as in this work. During this study, we created the following FE models: a) No-ACL model, that represents the injured joint, b) Native-ACL model, which represents the healthy knee, c) the ACLR - SB model that corresponds to standalone ACLR surgery and, d) the combined ACLR - LET model. The developed models are presented in Fig 2.

**Fig 2.**
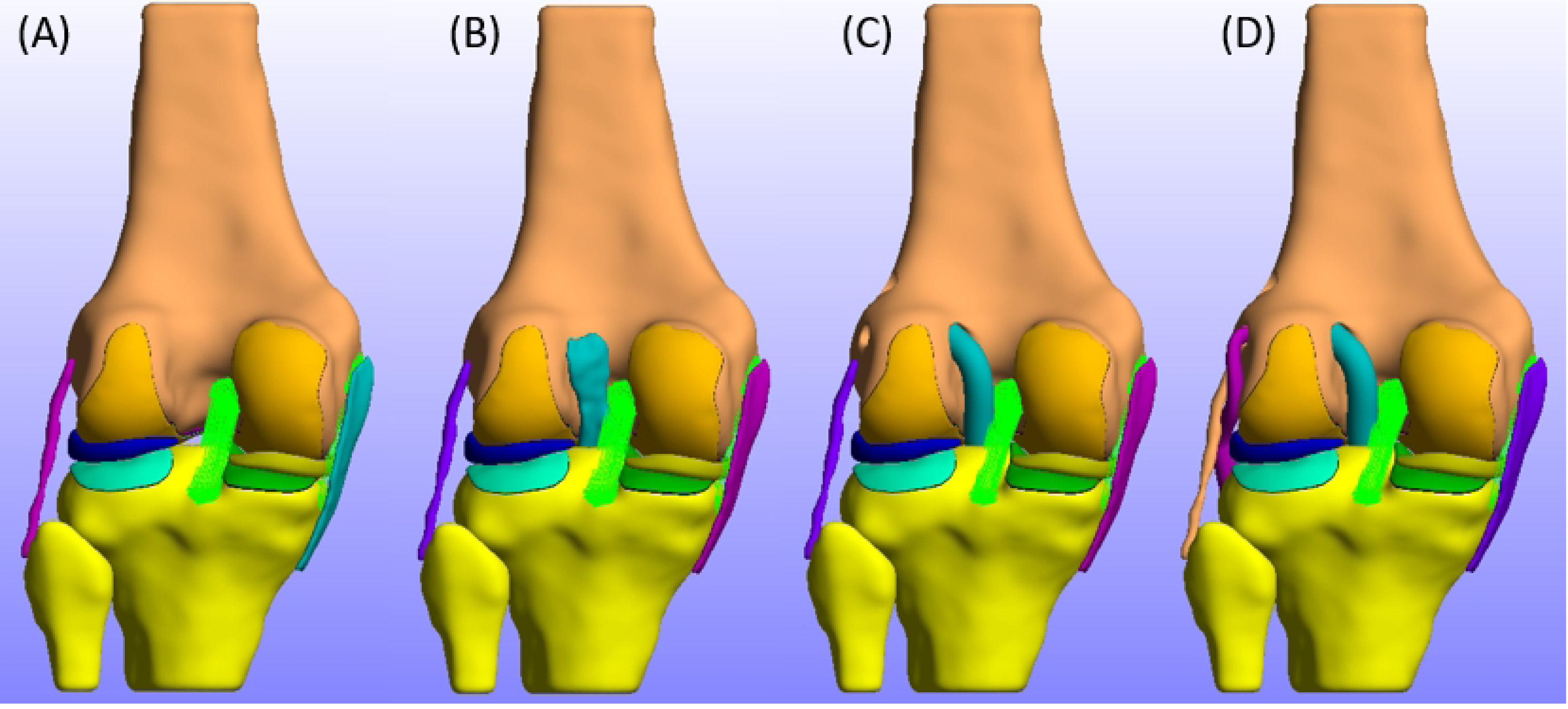
Developed FE models. In this figure we present form left to right the developed FE models used throughout this study. A: ACL - deficient knee FE model. B: Native = ACL FE model. C: Standalone SB - ACLR FE model. D: Combined ACLR - LET FE model.

### Geometries and material Properties

In the scope of this study, we adjusted a validated FE model used in our previous work [30]. The bones geometries were acquired directly from the Openknee (s) database in the case of the healthy and injured knee joint FE model. In the case of standalone ACLR and combined ACLR - LET the bones were modified by applying the “Surgery Modeling” as described above. We have decided to model the bones with rigid materials, since their deformation is considered negligible compared to the other present anatomical structures. This offers the advantage of prescribing kinematics using rigid joints that simulate knee joint motion in agreement to other studies [27, 40, 51, 54]. Moreover, the model complexity is greatly reduced since each rigid body is represented by a surface mesh, thus leading to tractable total simulation time. The femoral and tibial cartilages, and the menisci are modeled as in our previous study [30].

We replaced the spring ligaments for ACL, Medial Collateral Ligament (MCL), and LCL with volumetric meshes, that were generated using the Tetgen software [63]. The material properties were defined based on the corresponding validated OpenKnee FE model. To enhance the validity of the model, we reproduced the anterior laxity experimental tests of the OpenKnee (s) project in a FE analysis and adjusted accordingly the pre - strain value of the ACL ligament. This *in situ* strain is evident in biological tissues *in vivo* and contributes to joint stability [64]. The Posterior Cruciate Ligament (PCL) ligament was modeled as nonlinear tension-only springs with a force - displacement curve that was calibrated and validated in our previous study [30].

The grafts were modeled with an appropriate hyperelastic transversely isotropic Mooney - Rivlin material featuring an uncoupled deviatoric and volumetric behavior. The arithmetic values for the model parameters were defined based on curve-fitting on experimental data. We extracted the latter from studies performing uniaxial tests on the quadruple semitendinosus tendon and iliotibial band tissues [65, 66]. These are commonly used as harvesting sites among orthopedic surgeons during ACLR and LET [14, 16, 18]. Furthermore, we performed a series of uniaxial simulations to decide the optimal number of elements (mesh independence test) and proper values for the model parameters, such as the Bulk’s modulus. We provide a detailed description for determining the material properties in the supplementary material.

### Boundary conditions

In this section, we describe the applied boundary conditions for each model. Starting from contact definition, the menisci are rigidly attached to the tibia bone by properly selecting specific nodes that are close to the meniscal horns. The tibial and femoral cartilages are both rigidly attached to the respective bones by selecting the nodes of the most interior mesh layers. A sliding - elastic contact algorithm is used to model the interaction between the menisci and the cartilages and the femoral - tibial contacts. We performed iterative experiments to fine-tune the algorithm’s tolerance settings in order to achieve numerical stability and reduce mesh penetration. The same contact algorithm was used to model the interaction between each graft and the bone geometries.

The knee joint is modeled using the commonly adopted Grood and Suntay convention [67]. This requires the definition of the femoral and tibial reference frames based on anatomical landmarks. The kinematics are defined as a combination of three cylindrical joints creating a four - link kinematic chain. The femur and tibia coordinate systems were adopted by the OpenKnee (s) project [46, 59]. Before proceeding with the specific loading scenarios, we should clarify without loss of generality that in all cases the tibia reference frame is fixed in all Degrees Of Freedom (DoFs). The knee flexion is prescribed as a rotation between the rigid bodies of the first cylindrical joint and the loads are applied to the femoral reference frame.

Regarding the PS loading scenarios, in the case of the No-ACL and the Native-ACL models the following simulation steps are defined: 1) Use the Pre - Strain FEBio plugin [64] to apply in-situ strain in the models, 2) Apply the PS profile, where the tibia rigid body is fixed in all DoFs and the femur is free in all directions.

Subsequently, in the case of ACLR - SB models, the simulation steps are: 1) Pre-Strain plugin application, 2) knee flexion up to the desired graft fixation angle, 3) application of graft pretension force at the free end of the graft and then, graft fixation (fix the free end nodes to the tibia rigid body reference), 4) knee full extension, 5) application of the PS profile.

Finally, for the ACLR - LET simulations, we additionally implement the following simulation steps after the ACLR graft fixation step: 1) knee flexion up to the LET graft fixation angle and, 2) LET graft pretension and fixation. It is a common practice to perform LET after ACLR [14, 68]. All other steps are the same. For all ACLR knee models, the tibia is fixed in all DoFs. The femur is fixed in the anterior - posterior and lateral - medial displacement DoFs and internal - external rotational Degree Of Freedom (DoF) during graft fixation and prior to PS loading, to avoid introduction of displacement or rotational bias to the model. When the PS profile is applied the femur is set free similar to the No-ACL and the Native-ACL simulations.

### Graft pretension

In this section, we describe the adopted methodology and boundary conditions for graft pretension and fixation. Knee flexion angle during graft fixation is an important aspect of ACLR. Therefore, the FE simulations begin with a prescribed knee flexion angle up to the desired graft fixation angle. This flexion angle was set to 25*^◦^* for ACLR [22] and 30*^◦^* for LET graft fixation [17, 18, 57].

Regarding graft pretension we start by estimating the average orientation of the osseous tunnel where each graft is going to be pulled through. This is the tibial tunnel in the case of SB ACLR surgery and the femoral tunnel for LET, respectively. The orientation is specified as the normal vector of the plane that is defined using the location of three points around each tunnel’s exit expressed at the global reference frame. Then, a forward simulation is performed where a knee flexion is prescribed using a ramp function in the range of 0*^◦^* - 90*^◦^* with a step of 10*^◦^*. The femur is free to move, whereas the tibia is fixed at all DoF. At each step, we extract the quaternion that describes the femoral rotation. Using Spherical Linear Interpolation (SLERP), we can extract the quaternion corresponding to any desired fixation angle and estimate the corresponding rotation matrix. This rotation matrix is then multiplied with the initial unit vector to approximate the direction of graft pretension force at the desired fixation angle. Then the direction vector is scaled by the desired pretension force magnitude. The tension is applied using a ramp function. A rigorous description of the process is provided in the supplementary material to ease reproducibility.

At the end of the graft pretension step, a specific set of nodes that are inside the tunnel are connected to the corresponding bone mesh, signaling the graft fixation phase. In the subsequent steps, the displacement of these nodes is identical to the displacement of the rigid bone material.

### Clinical pivot shift

In this section, we describe the boundary conditions that we applied to the FE models so as to simulate the knee kinematics during the clinical exam of PS. In the clinical setup, the knee is flexed at between 10*^◦^* - 20*^◦^* and a combined load of internal torque, valgus stress and usually anterior force are applied to the tibia. This is the subluxation phase. A test is marked as positive if an anterior subluxation of the lateral tibia plateau is apparent [69]. Subsequently, as the orthopedist flexes the knee a reduction of the loaded tibia occurs approximately between 20*^◦^* - 40*^◦^* [20–22]. This type of PS is also called the “reduction” test, since the PS test occurs during the knee flexion phase.

Although the specific results may differ across subjects, as a general consensus we can state that the knee kinematics during the application of an effective PS profile should exhibit two key features during the reduction phase: a) An abrupt anterior translation of the femur relative to the tibia, and b) a simultaneous excessive internal femoral rotation. These correspond to a posterior tibia translation and external rotation, respectively. There can be multiple PS profiles that may lead to the above described PS kinematics [20]. In this study, we adopted and modified a static loading profile from a robotic clinical study [22]. The applied profile is depicted in Fig 3. Starting from left to right and from top to bottom, we apply a posterior force of 25N on the femur up to an initial step of 10*^◦^* of knee flexion. For the rest of the simulation we apply an anterior force of 25N. Furthermore, we apply a varus torque of 7Nm and an internal torque of 5Nm. The anterior - posterior force, and varus and internal rotation torques are implemented using sigmoid functions. The transition area of each sigmoid corresponds to a knee flexion angle in the range of 20*^◦^* - 30*^◦^*. This is evident by observing that the monotonic abrupt change of each loading curve is specifically defined in the above range. Finally, we apply a compression force of 20N to bring the cartilages into contact and assist the FEBio solver. All forces are applied on the femur reference frame.

**Fig 3.**
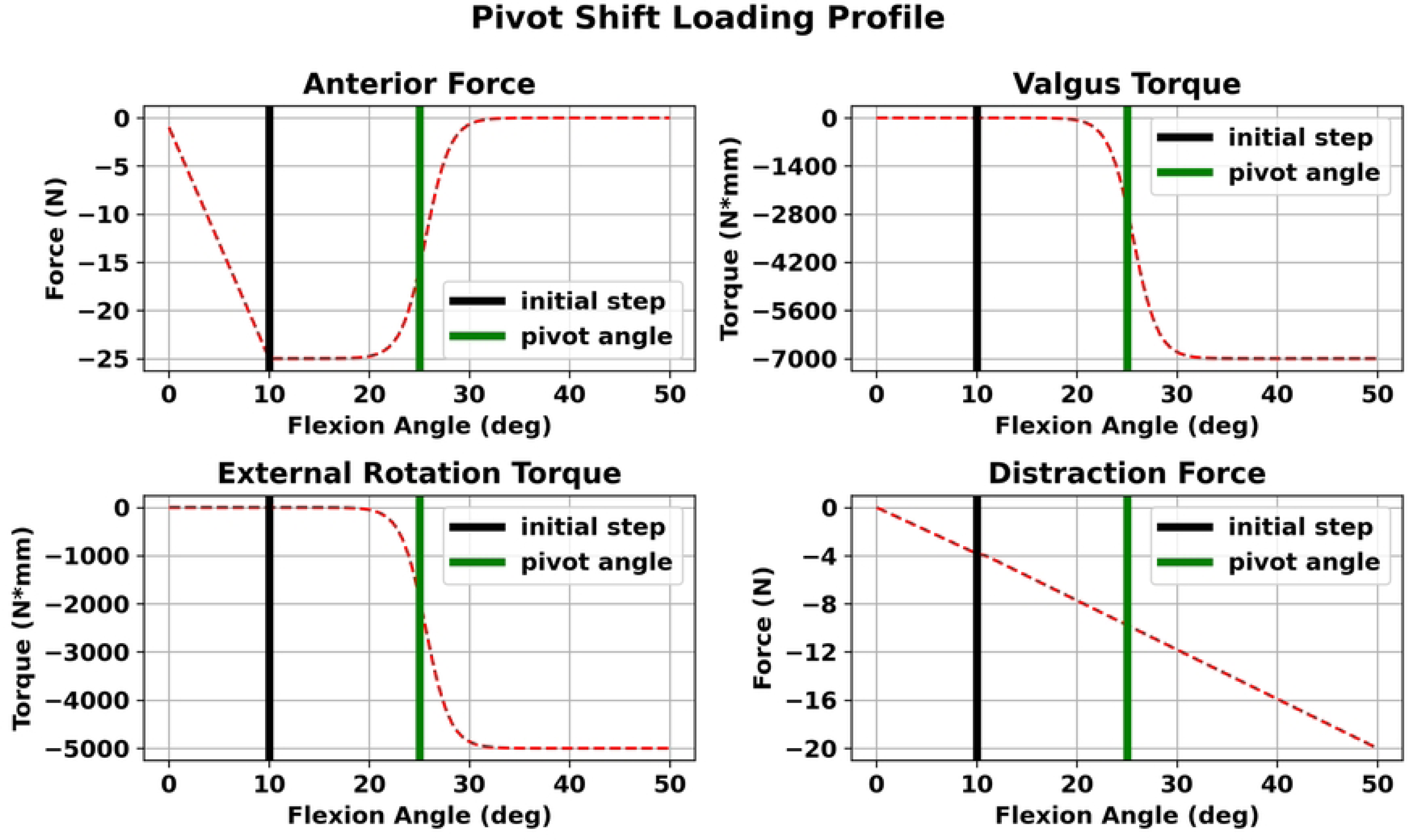
PS loading profile. Starting from the left to right and from the top to bottom, we observe the applied loads on the femoral reference frame to simulate the PS movement. A posterior force of 25N is applied up to an initial flexion angle of 10*^◦^* to induce the subluxation phase. As the knee flexes an anterior force is applied to model the reduction phase. Additionally, we apply a femur varus torque of 7 Nm and an internal femur moment of 5 Nm. These forces are implemented using sigmoid functions and are applied as the knee flexes from the initial angle of 10*^◦^*. The transition point of the sigmoids is set to 25*^◦^* (green vertical line). Finally, we apply a compression force to engage contact between the tibial and femoral cartilages and assist the FEBio solver. The profile is a modified version of the profile in [22].

### FE case studies

In advance of the results section, we think a comprehensive description of the FE case studies that we conducted in the scope of this work is imperative to assist the reader. Initially, we validated the behavior of the healthy ACL model by reproducing the anterior drawer test form the OpenKnee (s) project and adjusting the ligament’s *in situ* strain. Using FEBio’s pre-strain plugin this residual strain was applied by altering a fiber stretch to the ACL ligament material model. After each simulation run we estimated knee kinematics and compared them with the experimental ground truth data. We applied the Mean Squared Error (MSE) as our metric. Thus, the optimal fiber stretch was selected based on minimizing the MSE between the kinematics of each FE model and the experimental data.

Subsequently, we performed a FE simulation to assess the ability of the adopted PS scenario to produce the characteristic kinematic features of the clinical PS test. Towards this objective, we applied the profile to the No - ACL and the validated Native - ACL models and compared their performance.

Moreover, we compared the performance of standalone ACLR surgery and specific cases of ACLR - LET under the same PS profile and by applying different graft pretension values for both cases. We considered that the LET graft pretension is related to the degree of ETR reduction. Also, we wished to investigate whether the combined surgery provides better results for the same ACLR graft pretension as in the isolated surgery. Finally, the ACLR graft pretension is related to the developed graft stresses around the femoral tunnel. In both techniques the adopted tension values were found in literature. These correspond to 80N for the ACLR graft and 40N for the LET graft [16, 17, 23, 70, 71] The ACL graft tension was altered by a factor of *±*50%. The LET graft tension values ranged from 5N (“minimal tension”) up to +50 % of the literature value.

Finally, we performed a sensitivity analysis for assessing how different combinations of ACLR and LET graft pretension values affect the model’s performance in restraining the ETR and ATT. We wished to assess the correlation between different pretension values of both the grafts in reducing kinematic laxity and ACLR graft stress. Regarding ATT we adopted the following methodology. We identified the most lateral and medial aspects of the tibia plateau points. Their distance is the tibia plateau width. Then, the Tibia Medial Compartment (TMC) and Tibia Lateral Compartment (TLC) center points are estimated at 25 % and 75 % of the tibia width and along the line connecting the two aspects [22, 70]. Afterwards, we projected these points on the femur surface by estimating the vertices of the femur mesh that are closest from the line passing through each center point and with a direction along the z-axis. These point are presented in Fig 4.

**Fig 4.**
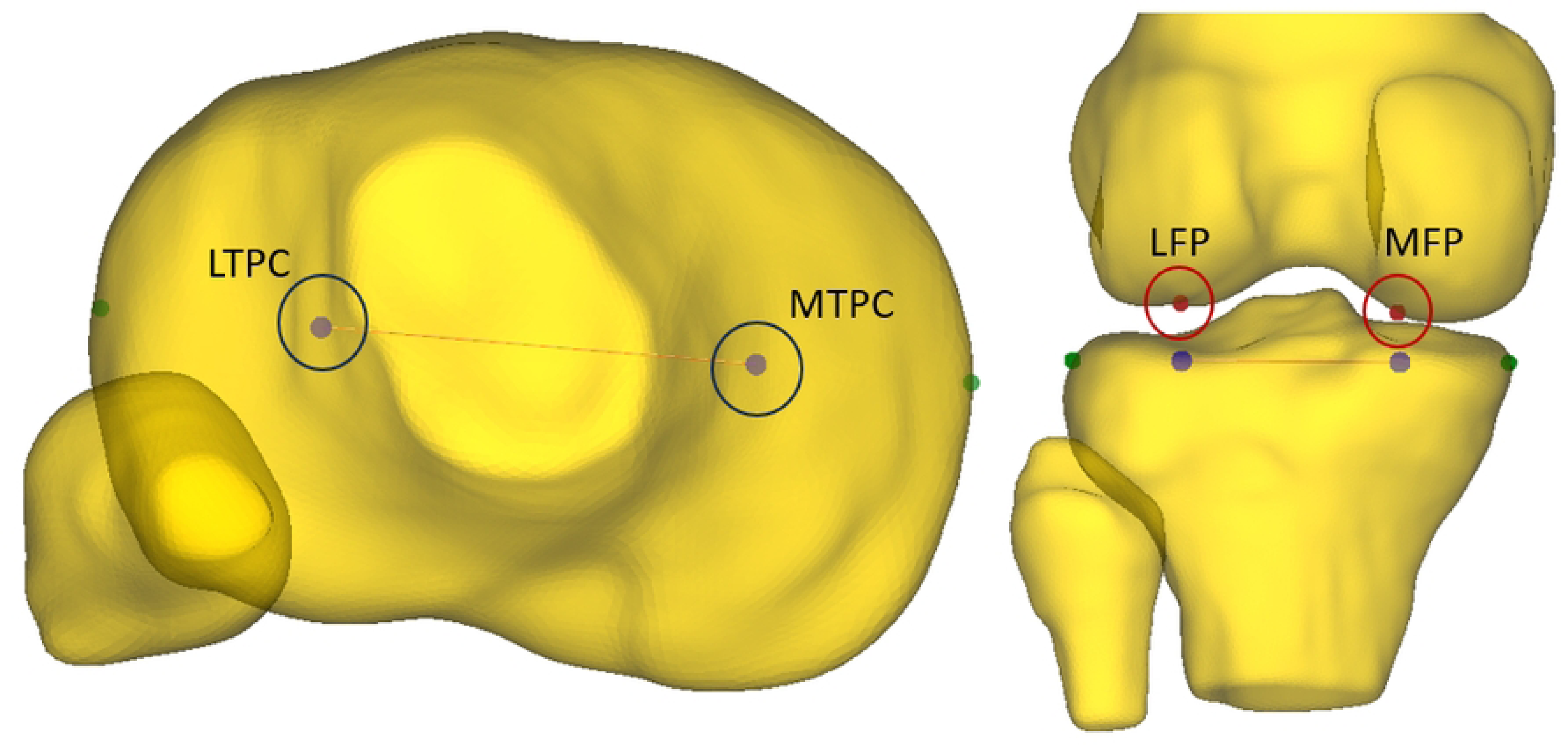
Femur projections of TLC and TMC Points. In this figure, we illustrate the estimated TLC and TMC points and their projections on the femur mesh. These points are taken at 25% and 75% of the tibia width [70]. The TLC projection on the femur (LFP) is used to measure the ATT during the PS simulation.

In the following sections we refer to the translation of the TLC when assessing ATT. Last but no least, we also assessed the developed stress around the femoral tunnel since we have already gained confidence in our graft mesh resolution.

## Results

In this section, we present the results for each simulation case starting from the validation of the baseline FE model. Then, we proceed with the assessment of the adopted PS profile. Subsequently, we compare the performance of standalone ACLR versus ACLR - LET for specific graft pretension scenarios found in literature. Next, we present the results for all combinations of pretension between ACLR and LET grafts using contour plots. The variables of interest are ETR and ATT of the TLC. We provide the graphs illustrating the evolution of these variables during the PS simulation step for each case. We also present the magnitude of these variables during the PS movement in a bar plot to provide an intuitive comparison between the reconstructed knee models and the healthy knee joint.

### Native - ACL model validation

In Fig 5, we present the sensitivity analysis results for the pre-strain assigned to the ACL material model. This residual strain was applied by defining a proper fiber stretch along the material’s fiber direction. For this purpose, we used the the Native - ACL FE model that includes the intact ACL material. Since *in - situ* strain affects joint mechanics we envisioned that a proper pre - strain would enhance our model’s validity. Towards this direction, we used experimental data from the OpenKnee (s) project. These included a loading profile representing an anterior drawer test and the resulted knee kinematics. We focused on ATT and used linear regression to fit a line to the experimental values (red line in Fig 5). This line served as the baseline for assessing subsequent FE simulations where we reproduced the same experiment using the Native - ACL FE model as stated above. At each simulation we altered the fiber stretch applied to the ACL material model, we applied the same loading profile, and finally we estimated the ATT. The optimal fiber stretch corresponded to the model that exhibited the lowest MSE between the simulated ATT and the reference red line. In our case, this fiber stretch was found to be a value of 1.06. Therefore, we assembled a validated ACL model that could be used as a reference when assessing the performance of the ACLR FE.

**Fig 5.**
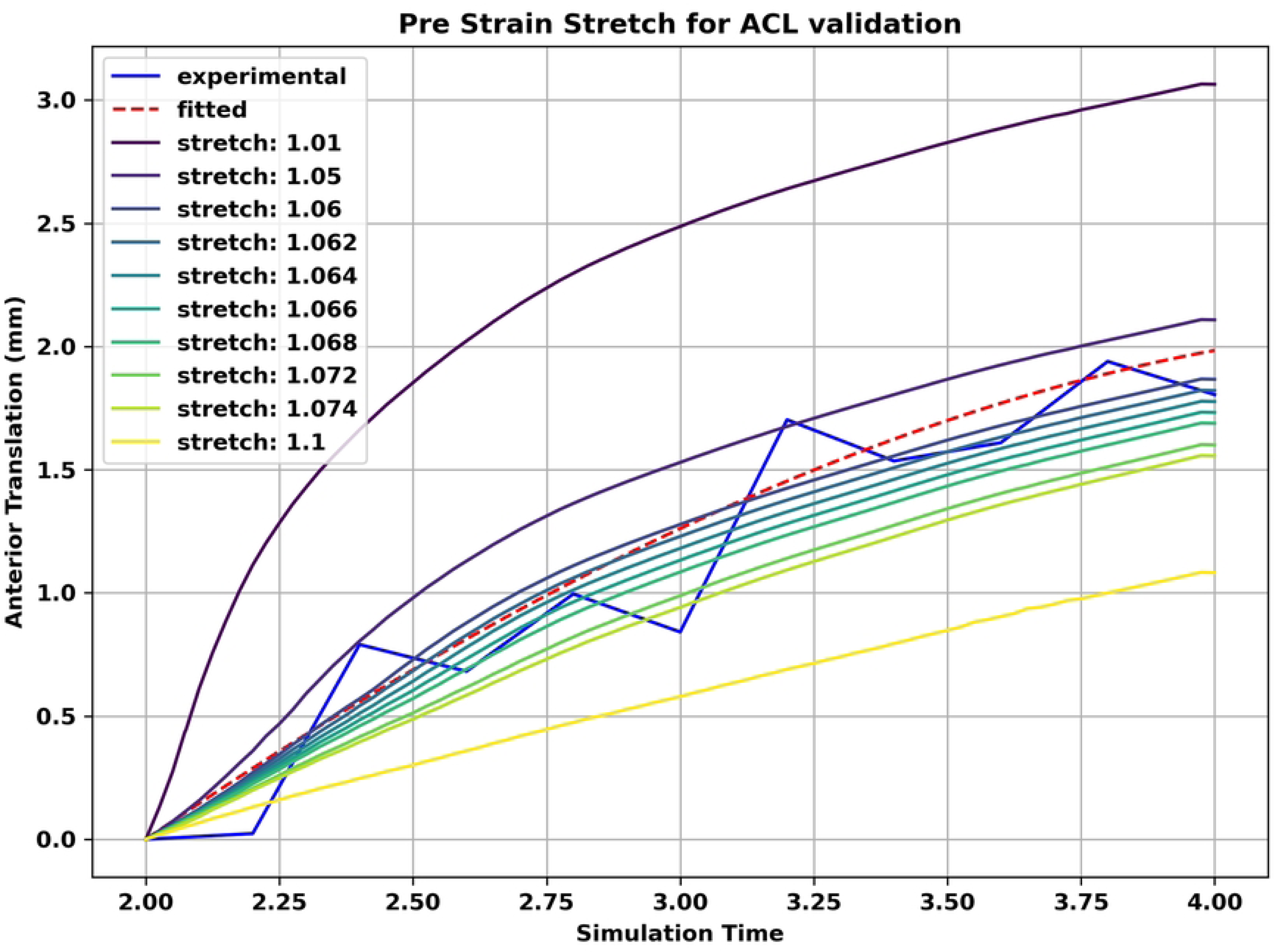
Validation of Native - ACL model regarding ACL pre - strain. In this figure, we present the results for deciding the optimal fiber stretch that was applied as *in situ* strain in the ACL material. Since this strain affects joint mechanics, we performed a series fo FE simulations to reproduce an anterior drawer experiment from the OpenKnee (s) project. During each simulation we altered the fiber stretch appointed to the ACL material, applied the experimental loads and measured ATT. We estimated the MSE between simulated ATT and the red reference line which was fitted to the experimental ATT (blue line). The lowest MSE was found for a fiber stretch of 1.06.

### PS profile assessment

In this section, we discuss the effectiveness of the adopted PS profile, which is presented in Fig 3. We applied the loading profile to the FE models representing the injured and healthy knee joint respectively and evaluated the ATT and ETR. The results are presented in Fig 6. We observe that the PS characteristic features are evident as depicted by the abrupt changes in both ETR and ATT. The PS movement occurs at a knee flexion angle of approximately 25*^◦^*. The applied loads cause the tibia in the ACL deficient knee (black dashed line) to start from an initially subluxed configuration that is characterized by the excessive ATT of the TLC and an increased internal rotation. The tibia is rapidly rotated externally and translated posteriorly highlighting the two prominent features of the clinical PS starting at approximately 23*^◦^* of knee flexion angle. The ETR value is 19.94*^◦^* and the respective ATT is -16.89 mm. We also observe similar behavior for the Native - ACL. However the magnitude for both variables is much lower with an ETR of 8.159*^◦^* and an ATT of -4.656 mm.

**Fig 6.**
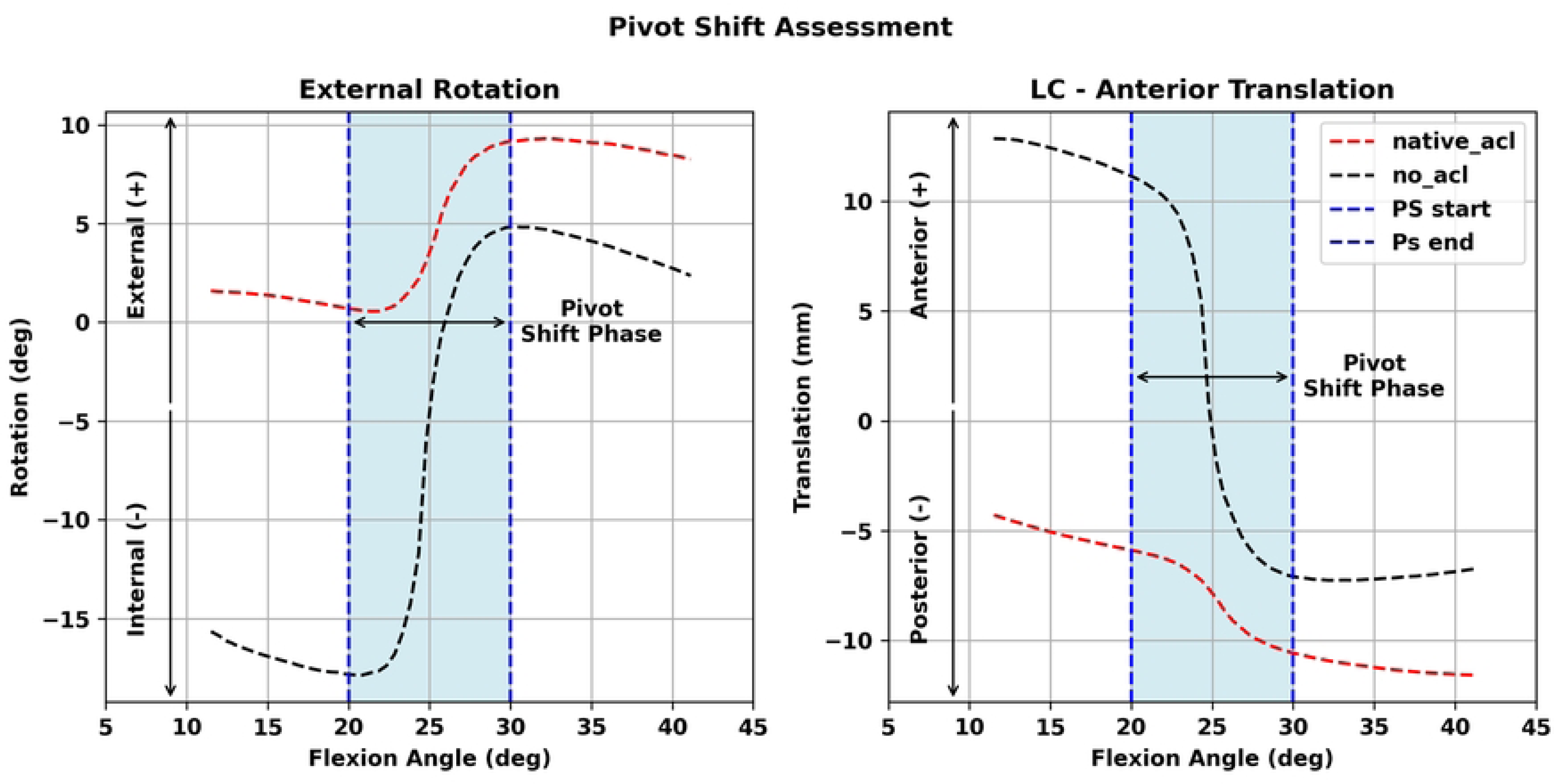
PS loading profile assessment. In this figure, we illustrate the effect of the applied PS loading profile on the ETR and ATT for both the No - ACL and Native - ACL cases, respectively. The PS movement is clearly evident at about 25*^◦^* of knee flexion with an abrupt ETR and posterior translation (negative ATT), respectively. The PS phase is highlighted with light blue color and spans the range between 20*^◦^* - 30*^◦^*. The adopted PS is qualitatively similar to clinical and simulated PS profiles [20, 72]. (LC): TLC, (PS): Pivot Shift.

Comparing these results with clinical studies that performed PS tests using robot simulators, we can find values in the range of (-0.5, -5.0) mm for ATT and *−*0.01*^◦^* - 3.0*^◦^* for ETR of the Native - ACL knee [20]. In the same study, a clinically performed PS demonstrated an ATT of -10 mm and an ETR of approximately 15*^◦^*. Loading profiles from this study were also adopted by other clinical trials [68]. Other studies reported values of -8 up to -12 mm of ATT and 15*^◦^* - 21*^◦^* of tibia rotation [72]. Similarly, a simulated PS in another clinical study showcased an anterolateral subluxation of 12.5 mm at a flexion angle of 30*^◦^* [58]. In an other clinical study, 25 examiners performed PS test on two cadaver knees with different degree of rotational laxity [73]. The results for the deficient knee demonstrated an average ATT of 8.8 mm in the low - laxity knee and an average of 17.3 mm in the high - laxity joint. The respective values for ETR were 10.0*^◦^* and 19.3*^◦^*. The values for the high - laxity knee are almost identical with our findings.

The difference in ATT can be associated with the difference in applied anterior force during the PS. Likewise, discrepancies in ETR can be related to differences in external torque applied to the tibia. Nonetheless, our PS profile exhibits similar qualitative behavior and comparable values for both ATT and ETR. Therefore, we can claim that the adopted profile is capable of producing a PS simulation setup to assess the performance of ACLR surgery techniques.

### Standalone ACLR

In this section, we present the performance of the standalone SB ACLR surgery during the PS simulation. We varied the ACLR graft pretension the range of 80*N ±* 50%. The results are presented in Fig7. The top charts present the evolution of ETR and ATT per knee flexion angle. At the bottom we provide bar plots that describe the magnitude of each variable based on their range during thePS phase. We observe that increasing the graft tension leads to reduced ETR and ATT. However, the ETR is larger when compared to the Native - ACL model as seen from the bar plot in the lower left part of Fig7. This holds true for all cases of graft pretension values. We notice the same behavior for the ATT of the TLC where the increase of the graft tension leads to a decrease of ATT and closer to the Native - ACL model. However, we can notice that pretension of the ACLR graft does not have a great effect on these two variables.

**Fig 7.**
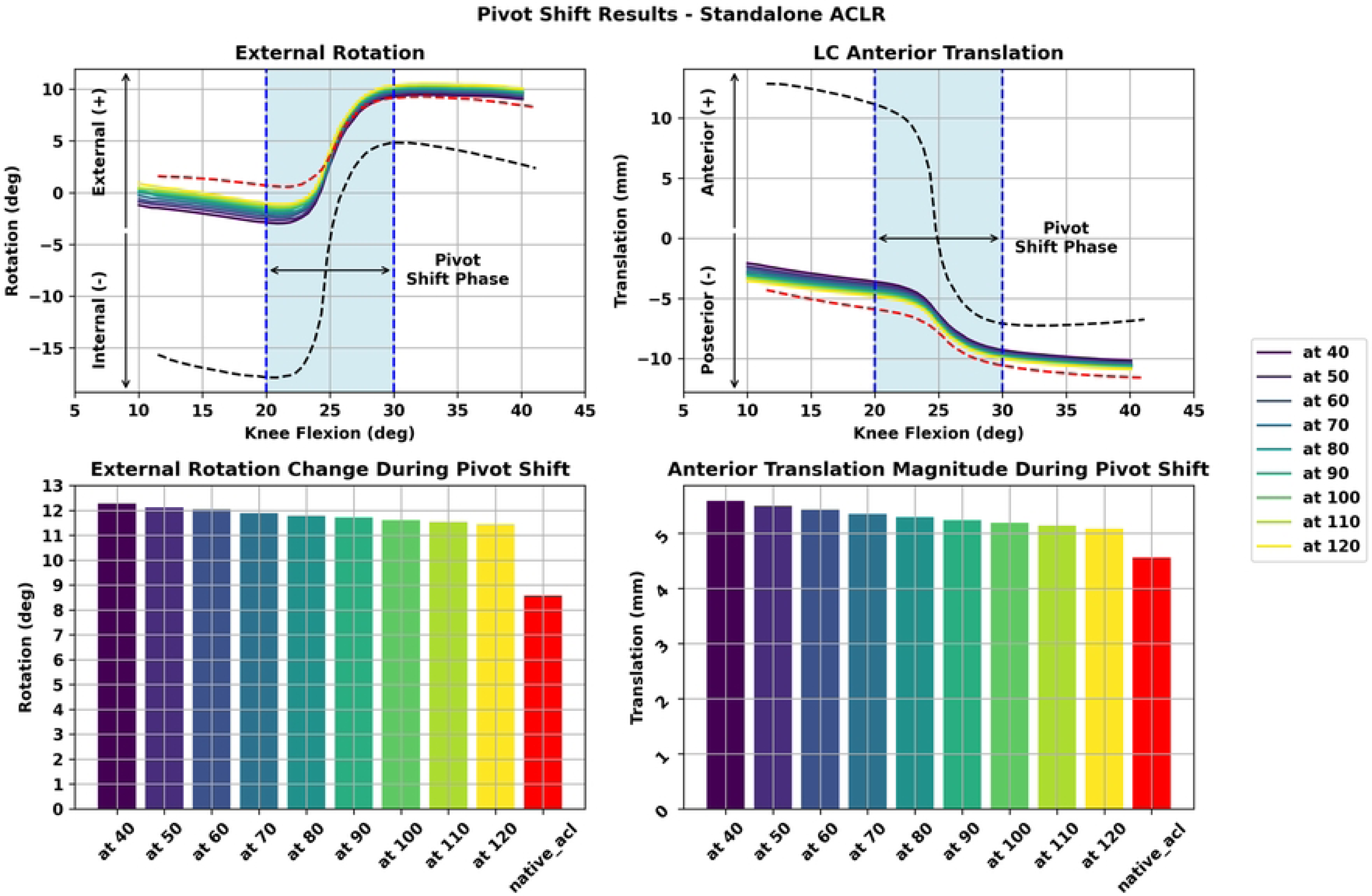
Assessment of standalone ACLR during PS. In this figure, we illustrate the performance of standalone ACLR in restoring the native tissue ETR and ATT. At the bottom we present the magnitude of both variables ranges during the PS movement using bar plots. As the graft pretension increases both variables reduce. In both cases, the ACLR knee model exhibits greater ETR and ATT values compared to the Native - ACL model. **(at)**: ACLR tension.

### Combined ACLR - LET

In this section, we illustrate the performance of the combined ACLR - LET surgery. We modified the LET graft tension while the ACLR graft tension was fixed in a certain value (80N). The results are depicted in Fig 8.

**Fig 8.**
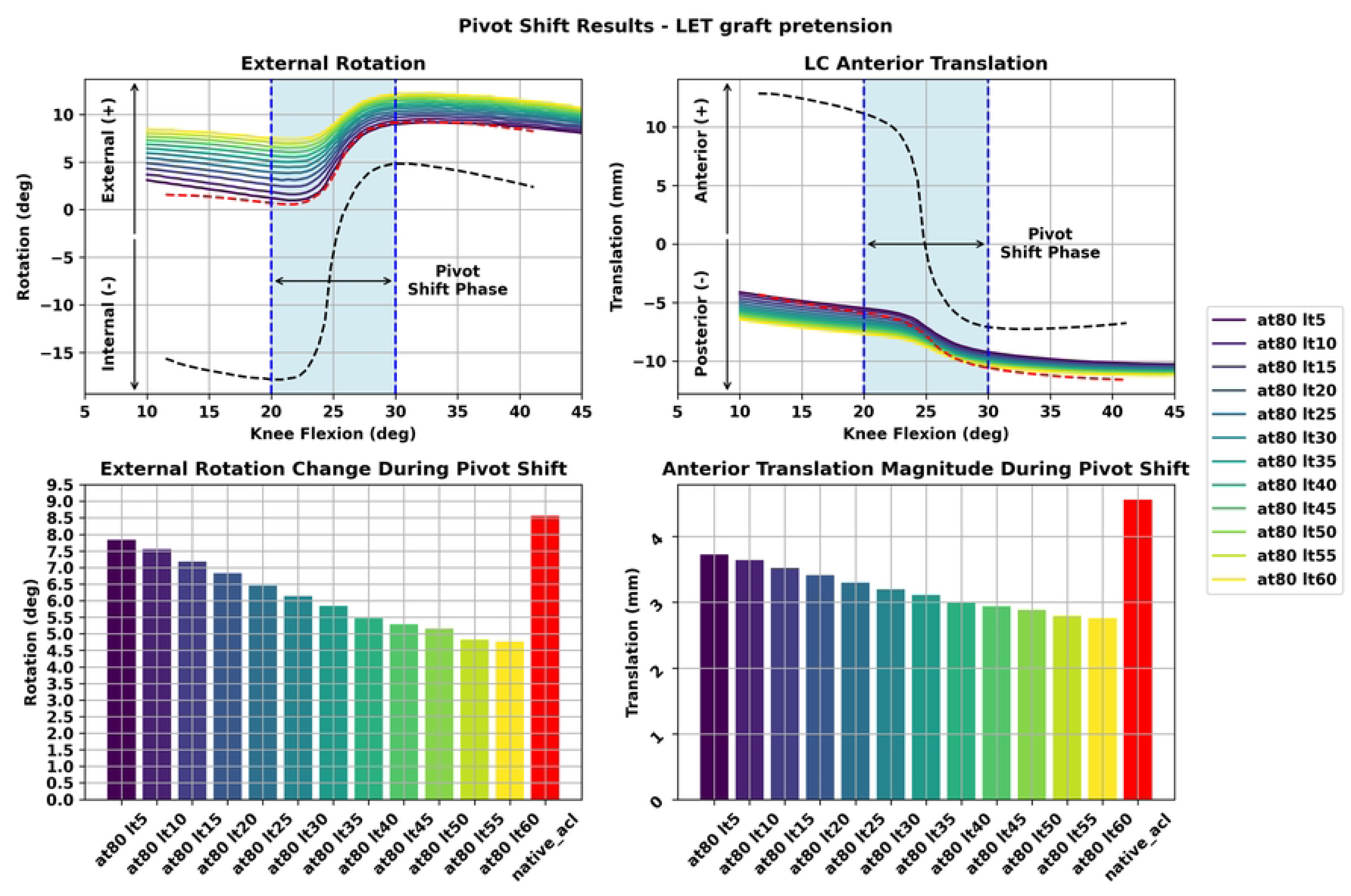
Assessment of LET graft pretension for the combined ACLR - LET during PS. In this figure, we depict the results for the scenario where the LET graft pretension is altered and the ACL graft tension is kept constant. We observe that increasing LET graft pretension leads to decreased ETR and ATT. However, in all cases these values are lower than the performance of the Native - ACL model. **(at)**: ACLR tension. **(lt)**: LET tension.

We observe that increasing the graft pretension load leads to reduced ETR and ATT. However, in all cases the achieved ETR is lower than the performance of the Native - ACL model and with a downward trend. The same holds true for the ATT variable as seen from the bar plot on the lower right of Fig 8. Therefore, we can state the increasing the LET graft pretension leads to a decrease in both ETR and ATT. However, larger values of tension may lead to excessive knee constraint for both the ETR and ATT variables.

### Pretension sensitivity analysis for combined ACLR - LET

Subsequently, we decided to identify the relationship between the pretension values for the ACLR and LET grafts, respectively, in the scope of the combined ACLR - LET surgery setup. The objective was to find the optimal combination that leads to reduced but no excessive rotational and translational laxity. Therefore, we analyzed 108 FE models that correspond to all possible combinations of graft tension values for the ACLR and LET grafts. We mention again that the ACL tension values varied in the range of 80*N ±* 50% with a step of 5N. The respective LET graft tensions were between 5N - 60N with a step of 5N. We decided to depict the results using contour lines for both the ETR and ATT of the TLC. For both variables, our metric was the Mean Absolute Error (MAE) between each combination and the Native - ACL model.

The contour lines are presented in Fig 9. On the left we present the ETR and on the right the ATT results. Each combination is represented by a blue dot to accentuate its performance. Each colored area contains a range of values for both ETR and ATT that correspond to the MAE of these parameters during the PS movement. With yellow color we denote the lowest values and with purple the largest. Thus, the combinations of ACLR and LET graft tensions that belong to these yellow areas demonstrate the closest performance compared to the healthy knee FE mode during the PS phase. We also have denoted with a red star the combination featuring the lowest MAE for each variable. In the case of ETR this is the combination of 80N and 5N for ACLR and LET graft tension, respectively. The corresponding values for ATT are 110N and 10N.

**Fig 9.**
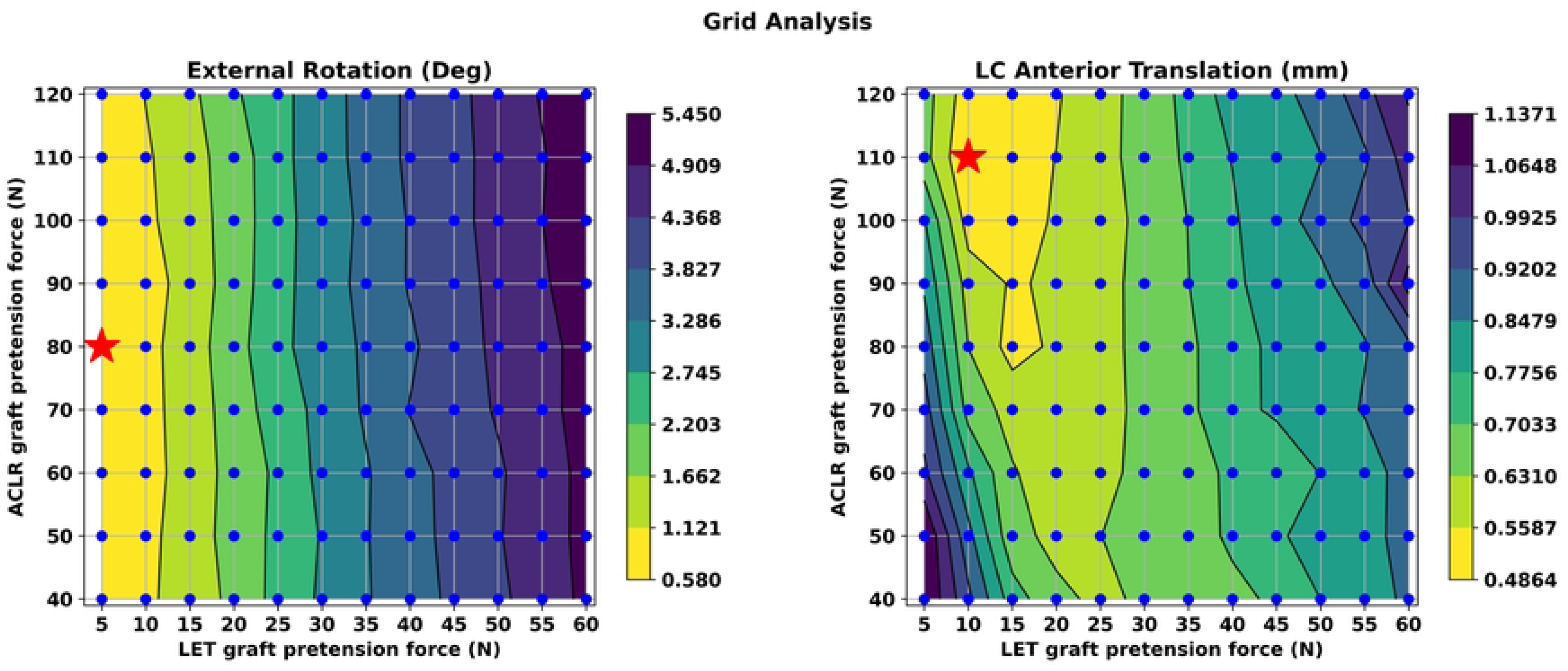
Grid analysis for ACLR and LET graft pretension. In this figure, we present the sensitivity analysis results for all combinations (ACLR tension, LET tension) between ACLR and LET graft pretension values using contour lines. These lines delineate different values of MAE between each combination and the Native - ACL model. The performance of each model is presented by a blue dot. Yellow areas demonstrate the smallest errors whereas purple areas the largest. We also highlight the best combination in terms of minimum MAE with a red star for both variables. These are the pairs (80, 5) and (110, 10) for ETR and ATT, respectively. We observe that ETR is more sensitive to LET tension as illustrated by the almost vertical alignment of the contour plot lines. On the other hand, larger values for both ACLR and LET grafts are required to reduce MAE inATT.

A general remark is that the ETR is more sensitive to the LET graft pretension value as depicted by the almost vertical alignment of the contour lines. Also, the MAE is reduced when we apply a smaller pretension value for the LET graft. On the contrary, we observe that the ATT demonstrates a different, very interesting non - linear behavior. We observe that the combinations with lowest MAE are in the range between 10N and 20N of LET graft pretension and from 80N up to 120N regarding ACLR graft pretension. Hence, larger values of pretension for both grafts are required to achieve an ATT that is comparable to the Native - ACL model.

### ACLR graft stress analysis

In Table 1, we present the ten best combinations between ACLR and LET graft tensions for the combined ACLR - LET surgery in terms of restoring ETR. The error column contains the MAE in ETR during the PS phase between each reconstructed FE model and the Native - ACL FE model starting from the lowest value. For each combination, we also exhibit the corresponding ATT values. We also provide max Von Mises stress values for the ACLR graft around the femoral tunnel insertion site. In an effort to compare the combined method with standalone ACLR surgery we present on the right side of Table 1 the corresponding results for the standalone ACLR surgery.

**Table 1.** The ten best combinations of pretension for the ACLR and LET grafts in the combined ACLR-LET surgery. The term “best” refers to the optimal restoration of native ETR. For each pair, we also provide the corresponding TLC ATT and von Mises stress values around the femoral tunnel.

| Combined ACLR - LET |  |  |  | Standalone ACLR |  |  |  |
| --- | --- | --- | --- | --- | --- | --- | --- |
| ACLR-LET<br>Tension (N) | ETR<br>Error (deg) <sup>†</sup> | TLC ATT<br>Error (mm) <sup>†</sup> | Stress<br>(MPa) <sup>‡</sup> | ACLR<br>Tension (N) | ETR<br>Error (deg) <sup>†</sup> | TLC ATT<br>Error (mm) <sup>†</sup> | Stress<br>(MPa) <sup>‡</sup> |
| **80 - 5** | 0.65893 | 1.01682 | 21.73 | **120** | 1.18827 | 0.78104 | 26.47 |
| **60 - 5** | 0.69128 | 1.12798 | 20.02 | **110** | 1.20258 | 0.87648 | 25.77 |
| **70 - 5** | 0.70082 | 1.07536 | 21.08 | **100** | 1.21663 | 0.97247 | 25.54 |
| **110 - 5** | 0.70670 | 0.77238 | 23.38 | **90** | 1.23251 | 1.06695 | 25.14 |
| **90 - 5** | 0.71689 | 0.95706 | 22.33 | **80** | 1.19098 | 1.15642 | 24.76 |
| **120 - 5** | 0.74478 | 0.78228 | 23.83 | **70** | 1.27499 | 1.29906 | 24.33 |
| **100 - 5** | 0.76263 | 0.90079 | 22.89 | **60** | 1.35159 | 1.43652 | 23.79 |
| **40 - 5** | 0.82038 | 1.25727 | 18.44 | **50** | 1.43482 | 1.57485 | 23.31 |
| **90 - 10** | 0.82586 | 0.71219 | 21.49 | **40** | 1.58253 | 1.74180 | 22.48 |
| **50 - 5** | 0.84644 | 1.25800 | 19.45 | - | - | - | - |

<sup>†</sup> The ETR and TLC - ATT errors are measured as the MAE during the PS movement between each pair and the Native - ACL model.
<sup>‡</sup> The von Mises stress corresponds to the maximum Von Mises stress developed in the ACLR graft and around the femoral tunnel insertion area.

We observe that all these combinations include a minimal pretension value of 5N for the LET graft apart from the (90, 10) combination and feature a ETR MAE less than 1*^◦^*. On the other hand, the MAE for the ETR is larger in standalone ACLR and for the same ACLR graft pretension values. Also, it increases as the ACLR graft tension decreases. In general, the combined ACLR - LET surgery demonstrates a ETR closer to the Native - ACL FE model compared to the standalone ACLR surgery.

Regarding ATT we notice that in the isolated surgery the MAE increments as the ACLR graft pretension value is reduced. For the combined method, we notice that between the ten combinations of lowest ETR, the pair of 90N and 10N of pretension for ACLR and LET grafts demonstrates the lowest MAE for ATT. Also, larger values of ACLR graft pretension lead to the decline of MAE. Hence, we can state that the ETR stability is more affected by the LET graft pretension, whereas larger values of ACLR graft pretension improve ATT stability of the TLC ATT.

Furthermore, we perceive that in all cases the combined approach leads to reduced ACLR graft stresses close to the femoral insertion area and for the same ACLR graft pretension. Normally, the stress is reduced when we apply lower pretension values for the ACLR graft. For example, we notice that for the best combination regarding ETR the ACLR graft pretension is 80N and the MAE is 0.658 93*^◦^* and with a graft stress of 21.73 MPa. For the same pretension the isolated surgery leads to a ETR MAE of 1.188 27*^◦^* and a stress of 24.76 MPa. However, in the case of standalone ACLR the minimum MAE is for 120N of graft pretension that leads to increased graft stress at 26.47 MPa. In general, the differences between graft stresses are moderate. These results highlight that a trade-off between rotational and translational stability and ACLR graft stress should be addressed when planning ACLR surgery.

## Discussion

The essential objective of this study is to assess the effect of the combined ACLR - LET surgery approach on reducing knee ETR instability and restoring ATT employing a PS FE simulation study. Towards this direction, we deployed an automated modeling workflow that is built upon open-source software tools, and was previously published [30]. We can deploy this pipeline to create subject - specific FE versions of the healthy, injured and ACLR reconstructed knee joint. In the confines of this work, we first created a validated version of the healthy knee joint that served as a reference model. Subsequently, we created ACLR FE that represent the SB ACLR and the combined SB ACLR - LET surgerical techniques. We applied boundary conditions that simulate the PS clinical exam and assessed the performance of the surgery techniques in terms of ETR and TLC ATT. Several clinical studies have been conducted in an effort to assess whether LET conducted along standalone ACLR can lead to improved knee rotational stability after ACLR [14, 16, 18]. However, and to the best of our knowledge, this is the first research work that attempts to shed light in this topic undertaking a FE simulation approach.

Initially, we described the surgery modeling approach where we used the Blender Python API to model the the different aspects of ACLR surgery. The workflow was already presented in our previous study [30]. It requires bone geometries acquired from MRI data and can model a compendium of surgery parameters such as tunnel location, graft radius, graft tension, graft fixation angle and graft material. By proper selection of these parameters, the tool can be executed to model different surgery techniques, such as SB, DB and LET. In this work, we utilized the tool to create the bone and graft geometries for the standalone SB ACLR and the combined ACLR - LET surgeries.

Subsequently, we included the created geometries to a FE model assembly process that is materialized and automated through a custom Python module. This module creates FE models that can be solved by the FEBio solver. We have made this module publicly available [74]. FE modeling and simulations provide a valuable tool for studying biomechanics. However, FE models are susceptible to modeling assumptions and many studies initiate by creating a reference model that usually represent the healthy joint [31, 33, 34, 36]. Similarly, we attempted to create a validated FE model representing the healthy knee joint that would serve as the baseline for subsequent ACLR simulations. Regarding ligament representation, we decided to model the ACL, PCL, and MCL with volumetric meshes similar to other studies [53, 75]. We assigned to them suitable hyperelastic transversely isotropic material models [76]. The material properties were adopted from the OpenKnee (s) project database. The PCL was modeled using nonlinear springs to reduce complexity without sacrificing realism similar to other FE studies [25, 77]. The pre - strain and stiffness of the PCL were validated during our previous study using experimental data from the OpenKnee (s) project. We reproduced anterior drawer experiments to identify a suitable pre - strain value for the ACL material. We found that applying a fiber stretch of 1.06 using the FEBio’s pre - strain plugin minimizes the MSE between the FE simulation and the experimental ground truth data. On the other hand, we did not include in our model the patellofemoral joint and the muscles that wrap around the joint. Although these modeling choices seem to eliminate essential details of the physical counterpart, we can claim that our model representation is satisfying enough for the scope of our study. We are not experimenting with dynamic voluntary movements, such as gait. Rather, we assess our models by simulating a passive clinical exam, the PS. Moreover, similar choices can be found in FE literature [9, 27, 29, 31, 32, 40]. Regarding, knee joint kinematics we adopted tibia and femur reference frames that were optimized using experimental data from the OpenKnee (s) project. Thus, we can claim that we have a valid FE representation of the healthy knee joint.

The development of the validated FE model provided us a baseline to evaluate ensuing FE simulations and perform “what - if” scenarios with a strong confidence in our results. Apart from a validated model that is a surrogate representation of the physical component we had to also apply realistic boundary conditions. Thus, the first step, was to design a loading profile that closely resembles the forces and torques applied to the knee joint during the real world PS clinical exam. We considered that an effective PS profile should induce an abrupt posterior translation and external rotation of the tibia bone during a flexion angle in the range of 20*^◦^* - 40*^◦^* degrees [20]. The adopted profile was presented in the “Results” section and includes a femoral internal rotation of 5Nm and varus torque of 7Nm along with an anterior force of 25N at about 25*^◦^* of knee flexion. It can be described as a “reduction” PS test. In similar FE ACLR studies, the authors adopted a “subluxation” PS profile that included internal tibial and valgus torques. The knee flexion was fixed in certain angles (0*^◦^*, 30*^◦^* [37, 49, 50]. The values for internal rotation moment ranged from 4 Nm up to 7.5 Nm, whereas the valgus torques ranged from 6.9 Nm up to 10Nm [37, 50]. Anterior tibia forces of 103 N or 134 N were also included during the subluxation phase [37, 49, 50]. In other FE studies the PS was a combination of a knee flexion of 20*^◦^* and a tibia torsion about a vertical axis [40, 48]. Although, our PS profile acts on the reverse direction compared to the mentioned FE studies, the adopted values for the torques are similar. A difference is observed in the anterior force. However, we selected a force of 25N, which is close to the typical femur weight when the clinician raises the leg before applying the PS test [72]. It is also close to the 30N of other clinical LET studies that apply a simulated PS test [16, 20]. Additionally, our study is the first where the knee is set free during the flexion phase, without applying the PS test in a fixed angle. Nonetheless, our results demonstrate that the adopted PS profile clearly achieves the ‘reduction’ characteristics, which are the prominent features of the PS test.

After assessing the performance of our validated model with an effective PS, we proceeded with the assembly of FE models that represented the ACLR knee joint. Still, we had to to validate them in terms of material properties and mesh density. Hence, we used experimental data from uniaxial tests to acquire validated model properties for the selected grafts of the ITB [78] and hamstrings tendon [65]. Moreover, we aimed to provide results for von Mises stress around the femoral tunnel insertion region, similar to other FE studies [31–33, 36, 39, 40, 45, 48, 54]. Therefore, a mesh density test was crucial to enhance the fidelity of our simulation results. We performed sensitivity tests for the graft mesh where the mesh resolution was increased until the maximum von Mises stress change at the fixation site was not significantly larger than a threshold of 10 % . Given these validated properties, we created the ACLR FE knee models.

In the scope of our study, we sought to answer the key question of whether a complementary LET operation to the standard ACLR surgery leads to improved stability in terms of ETR and ATT and reduced stress in the ACLR graft. Therefore, we used our validated models in FE simulations of the PS clinical exam. We assessed the effect of graft pretension in restraining ETR for both cases. We used the range of each variable during the PS phase as a measurement. Regarding ETR, we observed that the standalone SB ACLR surgery exhibits a reduced ETR as the graft tension value increases. However, this ETR is larger than the Native - ACL model regardless of graft pretension with values ranging from 2.96*^◦^* up to approximately 3.5*^◦^*. The same remarks are evident when observing the graphs for ATT of the TLC. However, in these cases the absolute difference between the ACLR models and the Native - ACL are smaller than in ETR. These results imply that the standalone ACLR surgery could be susceptible to increased rotatory instability and ATT. Also, a larger value of graft tension is required to reduce ETR and ATT and align them with the Native - ACL model.

We compared these findings with the combined ACLR - LET technique. Initially, we assigned different values of graft pretension to the LET graft while the ACLR graft tension was kept fixed to a common literature value. We estimated again the range of ETR and ATT during the PS phase. We observed that the combined ACLR - LET exhibits lower values regarding both ETR and ATT compared to the isolated method as illustrated in Fig 8. The difference is glaring in the rotational stability where we observe values ranging from 4.481*^◦^* up to 8.002*^◦^* for the ETR. The baseline measurement was about 8.5*^◦^*. Moreover, we noticed that increasing the LET graft pretension leads to excessive decrease of the ETR and ATT. In fact, a level of ETR that is half the performance of the Native - ACL model is evident for the larger values of LET graft pretension.

As a general verdict we can state that the combined ACLR - LET leads to reduced ATT and ETR compared to the healthy reference model. This conclusion is similar to relevant clinical studies [13, 16, 79–81] where PS tests were used to assess knee kinematics restoration either by performing cadaver experiments or systematic reviews of clinical trials and studies on the same field. Even though these primary outcomes were quite promising, we assumed that a more comprehensive analysis would shed additional light on this domain. Thus, we attempted to identify which combination between the pretension values assigned to the grafts in the case of ACLR - LET surgery leads to knee kinematics that are similar to the Native - ACL model. This time, we adopted MAE as our metric. The almost vertical lines of the contour plots presented in Fig 9 showcase the sensitivity of the combined method to the LET graft pretension regarding ETR. Moreover, the best results appear on the far left of the contour and in regions were the FE models with minimal LET tension values of 5 and 10 N lie. In fact, the best combination is the pair of 80N and 5N of ACLR and LET tension, respectively. These results reveal that a minimal LET graft tension should be used to avoid over constraining the knee in terms of ETR. This result is similar to the methodology of clinical papers where minimal values of pretension in LET are capable of limiting the PS phenomenon without excessive constraint of the knee rotation [14, 82]. On the other hand, large pretension values lead to an overconstrained knee behavior regarding ETR [57].

On the contrary, we notice a different behavior regarding ATT, as depicted on the right side of Fig. 9. In this case, the combinations with ACLR graft tension ranging from 80N - 120N and LET pretension in the range of 10N - 20N exhibit the lowest MAE. The most efficient combination is the 120N and 10N for the ACLR and LET pretension, respectively. Hence, we notice that ATT requires a higher pretension force for both grafts to be comparable to the performance of the Native - ACL model. Furthermore, we notice that the combined method leads to lower ATT MAE compared to the standalone ACLR surgery and for the same ACLR graft pretension. However, these discrepancies are moderate in most cases. Similar results are reported in clinical studies [16, 83]. Nonetheless, graft pretension is a debated issue among clinicians as it is clear from the large variance between applied forces [18, 57, 81, 82]. Our findings suggest minimal graft pretension level, although subject specific characteristics, and the inherent assumptions of the simulation approach should not be neglected.

Additionally, a potential advantage of performing LET is the reduced stress developed in the ACLR graft leading to lower graft failure chances. We presented max von Mises stress for the ACLR graft elements around the femoral tunnel insertion area. We observed that for identical pretension values the combined method leads to a modest decrease in graft stress with a mean value of 2.93 Mpa. The decrease in graft stress seems to be larger for lower values of graft tension. The biggest decrease is 4.04 MPa and corresponds to the 40N of ACLR graft pretension. Graft stress reduction is reported also by similar clinical studies [16, 58]. LET appears to act in a protective way regarding the ACLR graft. Nonetheless, the grade at which the ACLR graft is offloaded is still unclear and depends on several parameters, such as graft fixation angle and graft pretension. Also, the choice of passing the LET graft under or superficially to LCL seems to affect ACLR graft stress reduction [14, 16]. Still, this stress reduction is a potential contributor to the reduced graft failure rates when a combined surgery is adopted [14, 16, 58].

Although our results are promising and in general agreement with relevant clinical research, our study does not come without limitations. From a modeling point of view, a natural consequence of FE simulations is that assumptions must be made when approximating biomedical problems. These include loading boundary conditions, contact models and foremost, material properties. We tried to validate each step of our modeling workflow using experimental data. Also, the ITB was not included in our MRI data and we rather modeled the ITB strand with as much as possible realistic geometric features. Furthermore, key structures such as the patella and surrounding muscles are omitted in our analysis. This is justifiable in the case of a passive clinical exam such as the PS. However, the ACL injuries are most common in demanding physical tasks. We anticipate that all these limitations can be interesting directions of future work that could further enlighten the different aspects of ACLR. We can materialize these prospective research objectives improving the already developed modeling tools. We would like to assess the impact of LET in future FE simulations that involve boundary conditions from rigid multibody dynamics resembling such activities. For example, the developed lateral tibiofemoral contact pressures after LET or the impact of ITB debilitation to knee loading due to graft harvesting are debatable topics that we would like to address [57, 68, 84, 85]. Moreover, in the scope of this work we developed a loading profile that seems to trigger the desired PS movement for the given knee model. We should apply the profile in more samples to obtain a more systematic assessment of the loading conditions and the applied surgery methods. Also, we wish to expand our analysis to include more ACLR surgery techniques, such as the Anterolateral Ligament Reconstruction (ALLR). Nonetheless, the results of this study exhibit a promising potential in assessing combined ACLR - LET in a FE simulation setting.

## Conclusion

Despite of these limitations, we propose a complete workflow of ACLR modeling and FE simulation that can be applied to investigate the effectiveness of different surgery techniques on restoring knee kinematics. The developed tool can be easily modified to enable modeling of such advanced surgery methods showcasing its promising potential. Also, we have created an open - source module for automatic assembly of FE models compatible with the FEBio solver. These FE models include the ACLR geometries and can be easily deployed to perform biomechanics analyses and answer key medical questions. Our main focus was the modeling and simulation of the combined ACLR - LET surgery in an effort to investigate its advantages and implications. As far as we know, this is the first attempt on this controversial clinical topic adopting a simulation strategy. Our results are on par with similar clinical studies. We provide all models included in this study to the research community along with scripts that reproduce our results to solidify research pellucidity. The FE models can be used and modified from fellow researchers to explore a variety of ACLR topics and provide feedback and improvements in our work. In conclusion, we envisage that the developed tools supported by the promising relevant clinical results can emerge to a valuable supporting framework available to clinicians and biomedical researchers for ACLR study in a pre - surgery planning setting.

## Supporting information

**S1 File.** Supporting Information related to this publication. (PDF)

## Author Contributions

**Conceptualization:** Konstantinos Risvas

**Methodology:** Konstantinos Risvas, Dimitar Stanev, Konstantinos Moustakas

**Software:** Konstantinos Risvas, Dimitar Stanev

**Visualization:** Konstantinos Risvas

**Supervision:** Dimitar Stanev, Konstantinos Moustakas

**Writing – original draft:** Konstantinos Risvas

**Writing – review & editing:** Konstantinos Risvas, Dimitar Stanev, Konstantinos Moustakas

**Resources:** Konstantinos Moustakas

